# Histone ChIP-Seq identifies differential enhancer usage during chondrogenesis as critical for defining cell-type specificity

**DOI:** 10.1101/727370

**Authors:** Kathleen Cheung, Matthew J. Barter, Julia Falk, Carole Proctor, Louise N. Reynard, David A. Young

**Affiliations:** Skeletal Research Group, Institute of Genetic Medicine, Newcastle University, Central Parkway, Newcastle upon Tyne, NE1 3BZ, UK; Bioinformatics Support Unit, Faculty of Medical Sciences, Newcastle University, Newcastle-upon-Tyne, NE2 4HH, UK

## Abstract

Epigenetic mechanisms are known to regulate gene expression during chondrogenesis. In this study, we have characterised the epigenome during *in vitro* differentiation of human mesenchymal stem cells (hMSCs) into chondrocytes. Chromatin immunoprecipitation followed by next-generation sequencing (ChIP-seq) was used to assess a range of N-terminal post-transcriptional modifications (marks) to histone H3 lysines (H3K4me3, H3K4me1, H3K27ac, H3K27me3 and H3K36me3) in both hMSCs and differentiated chondrocytes. Chromatin states were characterised using histone ChIP-seq and *cis*-regulatory elements were identified in chondrocytes. Chondrocyte enhancers were associated with chondrogenesis related gene ontology (GO) terms. *In silico* analysis and integration of DNA methylation data with chondrogenesis chromatin states revealed that enhancers marked by histone marks H3K4me1 and H3K27ac were de-methylated during *in vitro* chondrogenesis. Similarity analysis between hMSC and chondrocyte chromatin states defined in this study with epigenomes of cell-types defined by the Roadmap Epigenomics project revealed that enhancers are more distinct between cell-types compared to other chromatin states. Motif analysis revealed that the transcription factor SOX9 is enriched in chondrocyte enhancers. Luciferase reporter assays confirmed that chondrocyte enhancers characterised in this study exhibited enhancer activity which may be modulated by inducing DNA methylation and SOX9 overexpression. Altogether, these integrated data illustrate the cross-talk between different epigenetic mechanisms during chondrocyte differentiation.

**Summary:** Human mesenchymal stem cells are able to differentiate into chondrocytes, the cell type found in cartilage, making them an accessible system to study gene regulation during this process. Epigenetic mechanisms such as histone modifications and DNA methylation together with transcription factor binding play a role in activating and repressing gene expression. In this study, we investigated the genome-wide histone modification changes during chondrocyte differentiation. Integration of this data with DNA methylation and SOX9 transcription factor ChIP-seq revealed epigenetic changes at gene enhancer elements. Regions of the genome that transition from non-enhancers to enhancers in chondrocytes are enriched for SOX9 transcription factor binding sites. Luciferase reporter assays revealed that enhancer activity may be modulated by manipulating DNA methylation and SOX9 expression. This study has defined important regulatory elements in chondrocytes which could serve as targets for future mechanistic studies.

## Introduction

Chondrogenesis is the process of differentiation of mesenchymal progenitors into chondrocytes. Articular cartilage, present in synovial joints, comprises extracellular matrix secreted by chondrocytes and has an important function in aiding the mobility of joints. As the only cell type present in articular cartilage, chondrocytes are responsible for the homeostasis of cartilage.

Chondrogenesis is a multi-step tightly regulated process mediated by growth and transcription factors, with the SOX9 transcription factor instrumental to the progression of chondrogenic differentiation [1] although not initiation [2]. Gene expression during chondrogenesis is in part regulated by dynamic epigenetic mechanisms such as DNA methylation and histone modifications [3,4]. MicroRNAs (miRNAs) and long non-coding RNAs (lncRNAs) also play a role in chondrogenesis [5–7]. Genome-wide histone modification changes have been observed during *in vitro* differentiation of MSCs into chondrocytes [8]. As well as development, epigenetic mechanisms are also known to be involved in disease. *Cis*-regulatory elements such as gene enhancers have been shown to be disrupted in cartilage pathologies. Deletions in a distal regulatory region of the *SOX9* transcription factor gene and within the *SOX9* gene itself both lead to campomelic dysplasia in humans [9,10]. Mutations in enhancers of collagen genes are also associated with chondrodysplasias [11,12]. Osteoarthritis (OA), an age-related cartilage degenerative disease, has a strong genetic component and to date, the vast majority of polymorphisms that confer an increased risk are located in non-coding regions of the genome, including enhancers [13,14]. There is evidence that the OA phenotype may be linked to the re-activation of developmental pathways [15]. These studies demonstrate that epigenetic mechanisms regulate gene expression in numerous biological processes. However, how these mechanisms affect gene expression is not fully understood in cartilage development and disease.

Mesenchymal stem cells (MSCs) are able to differentiate into chondrocytes and have been used to study chondrogenesis *in vitro*. Tissue engineering solutions to cartilage repair include autologous chondrocyte implantation, cartilage autografts and injection of MSCs into the damaged site [16,17]. However, these methods are not widely used and complications can arise from their application. Further knowledge of the regulatory processes that control gene expression during chondrocyte development is required to develop and improve models for cartilage regeneration. Usage of *in vitro* models for human chondrogenesis is crucial for understanding the changes that occur during normal development of human cartilage.

In this study, histone ChIP-seq (H3K4me3, H3K4me1, H3K27ac, H3K27me3 and H3K36me3) was performed in a scaffold-free *in vitro* model of human MSC (hMSC) chondrogenesis [18]. Analysis of histone ChIP-seq data revealed that large scale chromatin state changes occur during chondrogenesis and chondrocytes acquire cell-type specific enhancers upon differentiation. Integration of chromatin states with genome-wide DNA methylation data demonstrated that de-methylated CpG sites are located within H3K27ac and H3K4me1 marked enhancers during chondrogenesis. Motif analysis revealed that chondrocyte enhancers contain *SOX9* binding motifs. Altogether, our study provides a comprehensive analysis into the global epigenetic changes during MSC chondrogenesis and highlights the role of enhancers in defining cell type specificity.

## Results

### Chromatin state changes during chondrogenesis

Bone marrow derived hMSCs were differentiated into chondrocytes over 14 days using an *in vitro* transwell model of chondrogenesis. This scaffold-free model produces a cartilaginous disc which expresses matrix components such as type II collagen and sulphated glycosaminoglycans. Chondrogenic genes such as *SOX9* have been shown to be induced during the differentiation of hMSCs using this established and reproducible model of chondrogenesis [6,18,19].

Histone modifications H3K4me3, H3K4me1, H3K27ac, H3K27me3 and H3K36me3 were assayed in hMSCs and differentiated chondrocytes (day 14) using ChIP-seq. These histone marks were selected to reflect a wide range of regulatory states. H3K4me3 commonly marks active promoters, H3K4me1 and H3K27ac are found at active enhancers, H3K36me3 are located at actively transcribed regions and H3K27me3 marks transcriptionally repressed regions. The genome-wide profiles of each histone mark were as expected; the density of each histone mark differs across the genome with the active promoter mark H3K4me3 showing a high density of peaks close to transcriptional start sites (TSS; Fig. 1A). Histone modifications are known to influence gene transcription; therefore, histone mark enrichments were correlated to expression levels of genes in hMSCs and differentiated chondrocytes (Fig. 1B; Supplementary table 1). As expected, histone marks typically associated with transcriptional activity were enriched in highly expressed genes (Fig. 1C), which in differentiated chondrocytes included *COL2A1* and *ACAN*, genes that encode the two major ECM components of articular cartilage. In contrast, the transcriptionally repressive mark H3K27me3 showed a greater enrichment in genes with low expression levels in differentiated chondrocytes, including the osteoblast transcription factor *RUNX2*. This demonstrates that the histone ChIP-seq generated in hMSCs and differentiated chondrocyte exhibit expected genome-wide profiles and gene expression associations.

**Figure 1.**
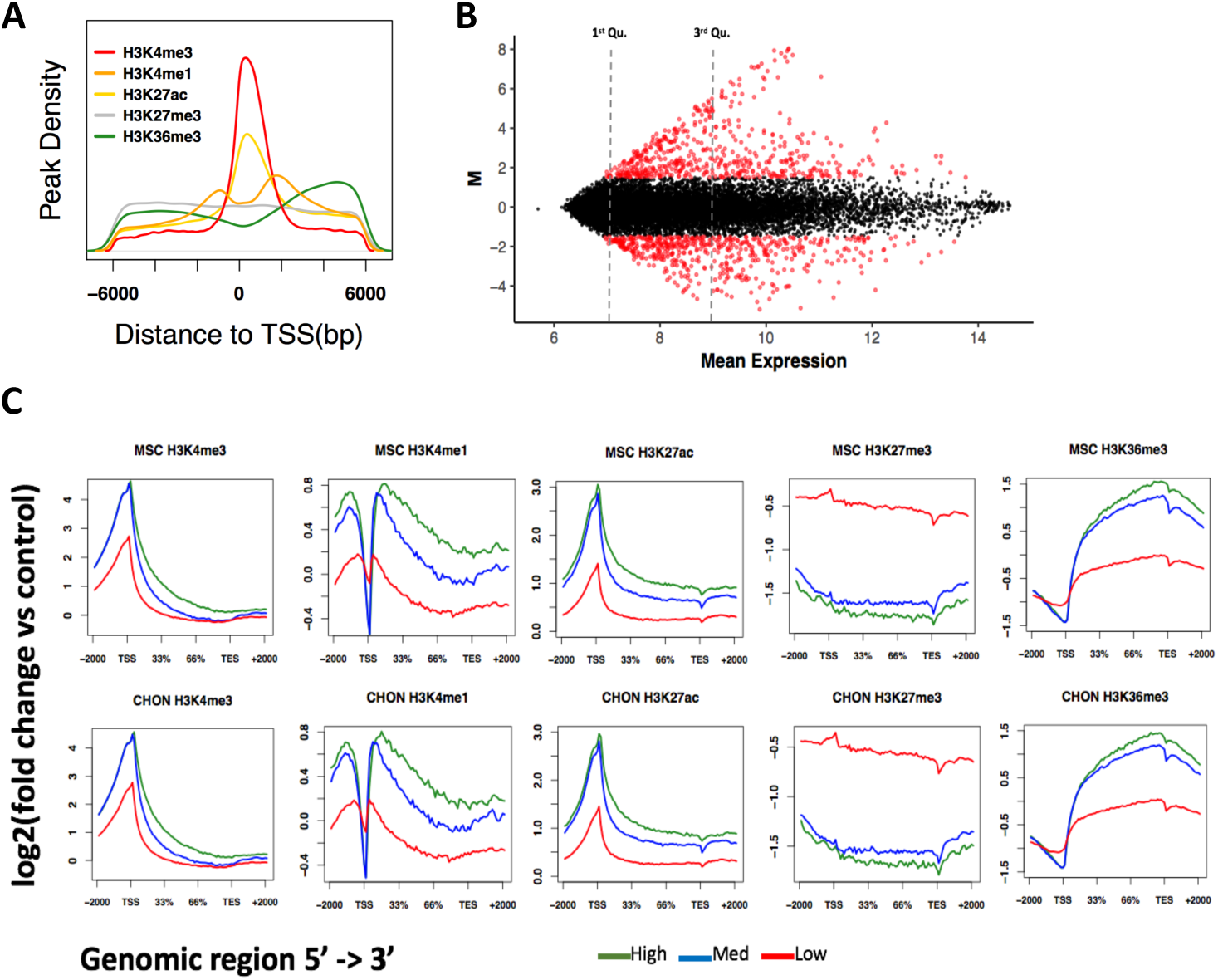
Correlation of histone mark enrichment with gene expression in hMSCs and differentiated chondrocytes. (A) Density of histone mark peaks around the TSS (+/-6kb). Peaks were called using MACS2 using input samples as controls for background noise. (B) MA (log ratio – average expression) plot of differentially expressed genes between hMSCs and differentiated chondrocytes. Gene expression was measured using a cDNA microarray and gene expression is reported as normalised signals. (C) Histone mark enrichments in genes categorised as have high expression (expression value > 9; upper quartile), medium expression (expression value > 7 and < 9) and low expression (expression value < 7; lower quartile). Histone marks were associated/overlapped to genes using the ngs.plot tool and plots were generated in R.

Combinations of histone modifications can reveal more information about the regulation of gene expression compared to singular histone marks [20]. Regulatory elements and chromatin states may be defined by the co-occurrences of specific histone marks [21]. A 16 chromatin state model was trained on the hMSC and differentiated chondrocyte ChIP-seq data using ChromHMM (Fig. 2A). The model yielded a range of chromatin states known to be associated with the histone modifications assayed in this study. This included promoter states, actively transcribed states and enhancer elements [22]. Large scale changes in chromatin states were observed between hMSCs and differentiated chondrocytes, particularly with regards to the quiescent and repressed states becoming transcriptionally active (Fig. 2B), demonstrating that genome-wide histone modification changes occur in the epigenome during chondrogenesis. To elucidate how chromatin states affect gene expression, the GREAT tool [23] was used to retrieve gene ontology (GO) terms for each chromatin state. GO terms associated with genes linked to each of the defined chromatin states were non-specific to cell-type and mostly encompassed general cell functions, the exception being enhancer states (Supplementary fig. S1-S15). In differentiated chondrocytes, the strong active enhancer state (characterised by high enrichments of H3K4me1 and H3K27ac; state 13_EnhS) yielded GO terms related to chondrogenesis and cartilage function (Fig. 2C). Previous studies have demonstrated that gene enhancers are cell-type specific and play an important role in regulating cell-type specific processes [24]. Accordingly, chondrocyte enhancers defined in this study are associated with chondrogenesis related terms, more so than promoter or gene transcription chromatin states. Chromatin state changes can clearly be observed around genes which show gene expression changes. For example, we observed the histone modification around the *COL2A1* gene switching from repressed/inactive in hMSCs to transcriptionally permissive in chondrocytes (Fig. 2D).

**Figure 2.**
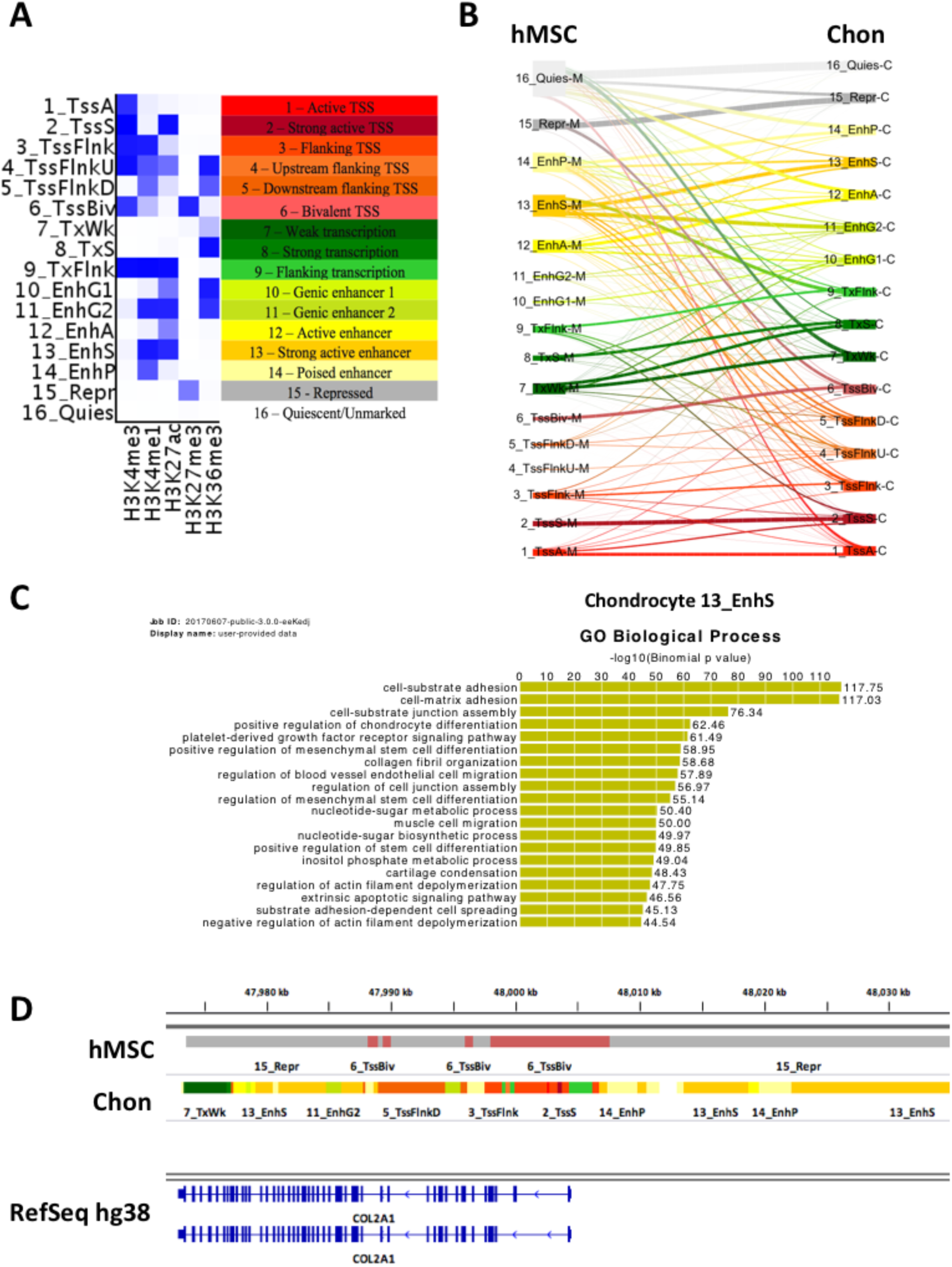
A 16 chromatin state model generated from hMSC and differentiated chondrocyte histone ChIP-seq. (A) Chromatin state model with annotated states. The chromatin state model was generated using the ChromHMM tool with all hMSC and differentiated chondrocyte data using input controls. (B) Genome wide changes in chromatin states during hMSC chondrogenesis. (C) GREAT biological process GO terms for the chondrocyte strong enhancer (13_EnhS) state. Genome co-ordinates of the 13_EnhS were used as input into the GREAT tool which associates cis-regulatory elements with genes. (D) IGV genome browser view of hMSC and chondrocyte chromatin states around the COL2A1 gene.

### Comparison to Roadmap Epigenomics cell types

Several large-scale consortia have aimed to characterise the epigenomes of various cell-types including the NIH Roadmap Epigenomics project [22] which defined chromatin states in 127 cell-types, 98 of which also included the active enhancer mark, H3K27ac. Roadmap cell-types contained bone marrow derived hMSCs and differentiated chondrocytes, therefore we sought to determine whether the epigenome of our chondrocytes were comparable to those included in the Roadmap project. We compared our 16 chromatin states to the equivalent states of the 18 state model generated by the Roadmap project for their 98 cell-types that contained the H3K27ac active enhancer mark (Supplementary fig. S16). The Jaccard similarity co-efficient was used to compare equivalent chromatin states across all cell-types in a pairwise manner. When individual chromatin states except enhancers, were investigated there appeared to be no apparent clustering of cells by type or origin (Supplementary fig. S17 – S22). In contrast, when H3K27ac and H3K4me1 marked enhancers (labelled 13_EnhS in transwell chondrogenesis chromatin state model and 9_EnhA1 in Roadmap 18 state model) were explored, cells clustered with other more closely related cell-types (Fig. 3). Our differentiated chondrocytes (“CHON” in Fig. 3) clustered together with the BM-MSC differentiated chondrocytes from the Roadmap project [8], demonstrating a higher level of similarity to each other that to all other cell types. The Roadmap bone marrow derived hMSCs and hMSCs in this study were closely related, contained within a small cluster of primary culture cells consisting of chondrocytes, myocytes, osteoblasts and fibroblasts (Fig. 3). These data corroborate previous studies which report that enhancers are distinct between cell-types, more so than any other regulatory features such as gene promoters [24,25]. Further, enhancers in chondrocytes from different sources showed higher similarity compared to other cell-type enhancers. Thus, there is a chondrocyte-specific epigenome based on gene enhancers that can be detected despite differences in chondrogenesis models, laboratory and MSCs donors.

**Figure 3.**
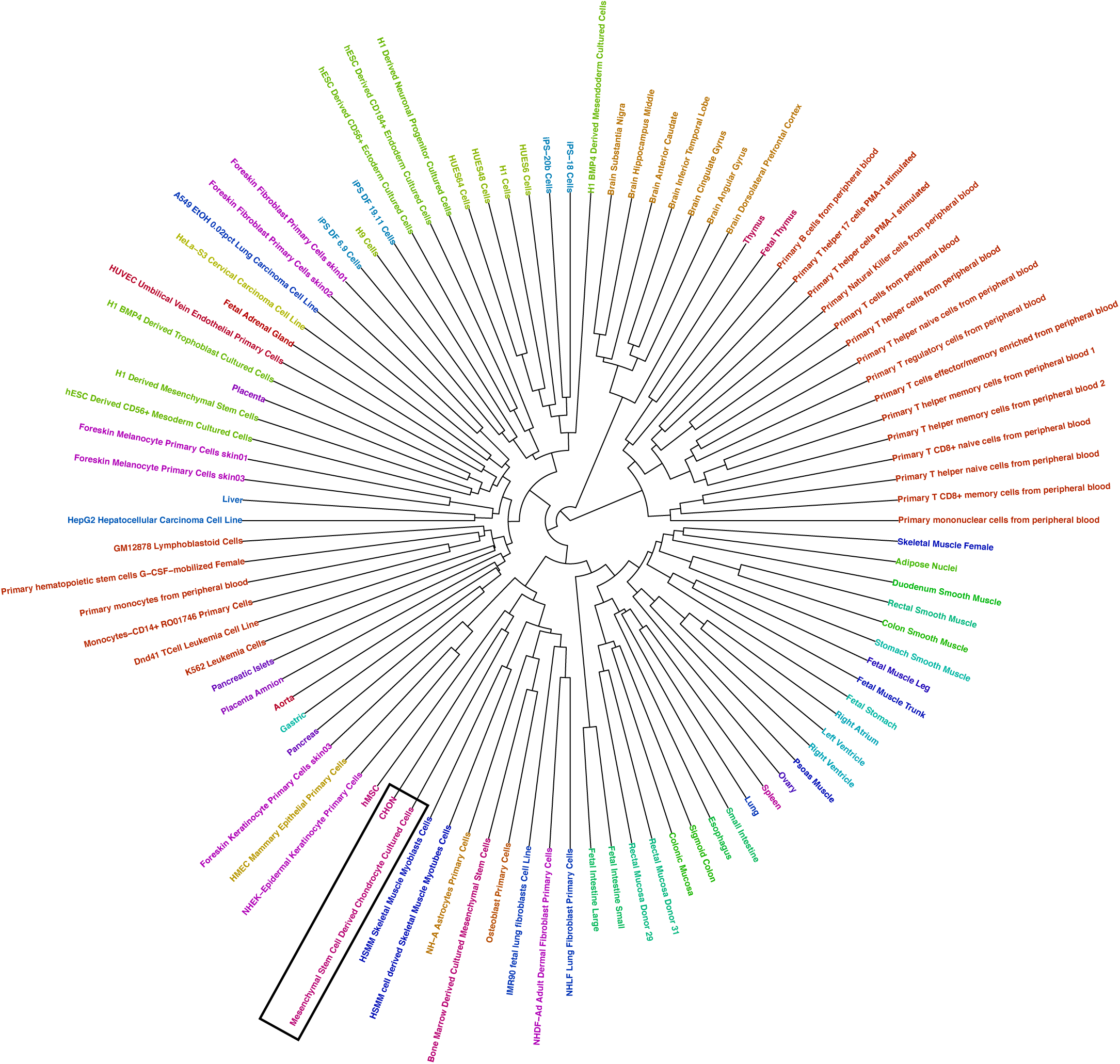
Clustered similarity heatmap of H3K4me1 and H3K27ac enhancers in hMSC and differentiated chondrocytes vs Roadmap cell types. The Jaccard similarity co-efficient was calculated between state 13_EnhS in the transwell chondrogenesis model and 9_EnhA1 in Roadmap 18 state model across cell types. Both states are characterised by high enrichment of H3K4me1 and H3K27ac. Values closer to 1 signify higher similarity. Stem cells and differentiated chondrocytes from this study are labelled “hMSC” and “CHON” respectively. Chondrocytes are indicated by the black boxes. A list of Roadmap cell types and their ID codes can be found on the Roadmap project website (http://www.roadmapepigenomics.org).

### DNA methylation at gene enhancers

Histone modifications are influenced by DNA methylation and vice versa during development [26]. DNA methylation occurs at CpG sites in the genome and is typically associated with transcriptional repression. An Illumina Infinium HumanMethylation450K BeadChip array was used to measure DNA methylation. DNA methylation changes during the *in vitro* transwell model of chondrogenesis were largely de-methylation events which were associated with chondrogenesis related GO terms [27]. We integrated the DNA methylation and ChIP-seq data in order to investigate the DNA methylation changes in chromatin states during MSC chondrogenesis, focusing on the hypomethylated CpGs (94% of the significantly differentially methylated loci during chondrogenesis) since this is linked to gene transcription activation. Global methylation patterns reflect known trends (Fig. 4A & 4B) e.g. gene promoters tend to have low percentage methylation relative to the rest of the genome [28]. We observed that enhancers marked by H3K4me1 and H3K27ac (13_EnhS state) were enriched for de-methylated CpG sites (Fig. 4C & Fig. 4D). Fewer than 2% of total CpGs probes present on the Infinium HumanMethylation450K BeadChip were located within chondrocyte chromatin state 13_EnhS (strong enhancers) yet remarkably 41.8% of de-methylated CpGs were found in this chromatin state (Fig. 4E), a highly significant over-representation (Chi-square test *p* < 0.001). We evaluated the effect of DNA methylation in six selected regions that acquired enhancer status during chondrogenesis in both our model and the chondrogenesis model in Roadmap and also overlapped with a H3K27ac signature during development in of human embryonic limbs [29]. The regions were cloned into a luciferase reporter vector with and without treatment of a CpG methyltransferase. Unmethylated enhancer regions showed increased enhancer activity compared to the empty vector control, confirming that regions classed as enhancers in our model exhibit enhancer activity. With the addition of a CpG methyltransferase, all regions showed a significant decrease in enhancer activity compared to unmethylated regions (Fig. 4F).

**Figure 4.**
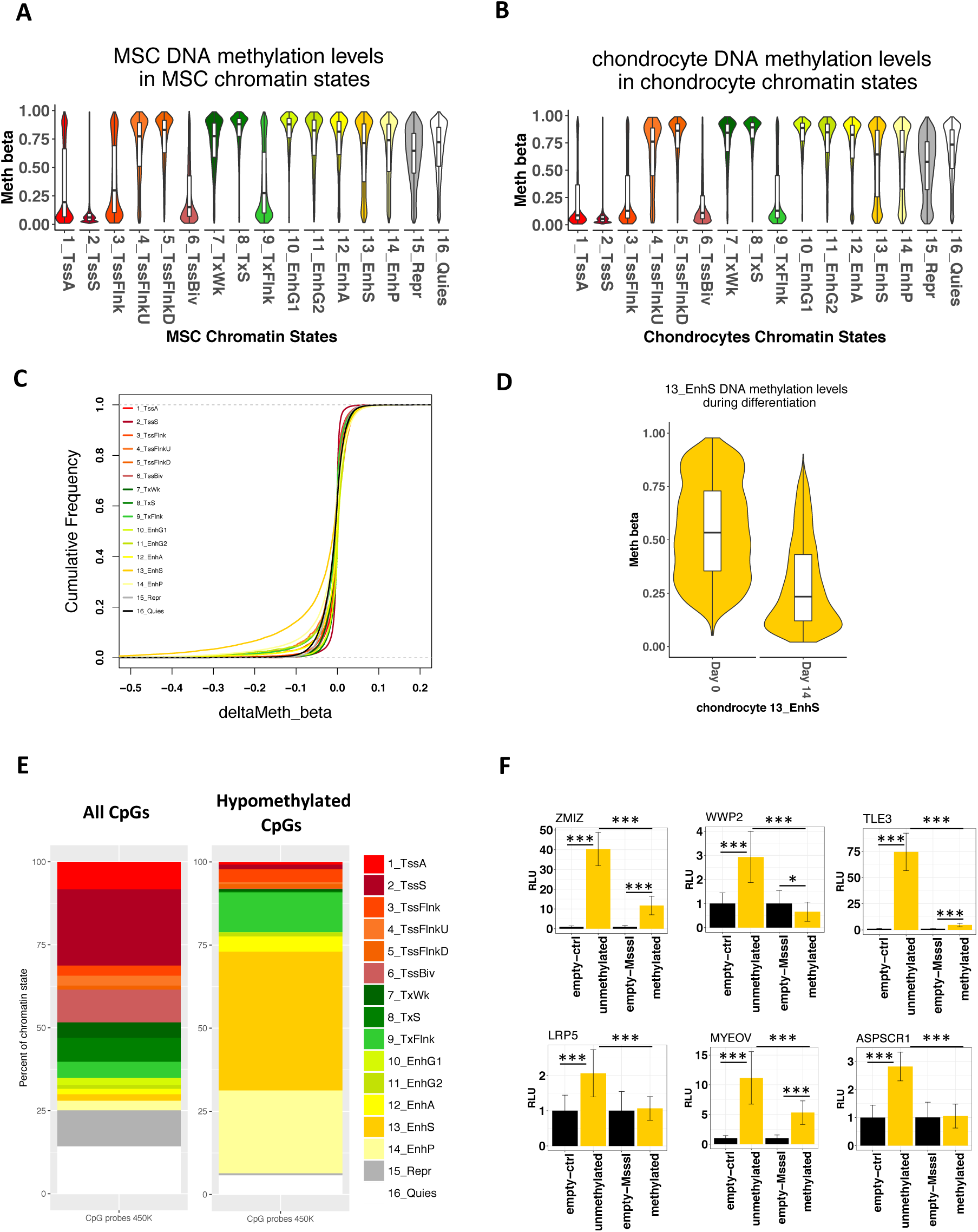
DNA methylation in chondrocyte chromatin states. (A) & (B) Methylation levels of CpGs in the hMSC and chondrocyte chromatin states respectively. CpG genome co-ordinates were intersected with chromatin states using BEDTools intersect. (C) Empirical cumulative frequency plot of methylation changes (beta values) in chondrocyte chromatin states during hMSC chondrogenesis. (D) Significantly methylated CpGs (FDR < 0.05) in between hMSCs and chondrocytes in chondrocyte chromatin states. (E) The percentage of all CpGs on the 450k array in each chondrocyte chromatin state and the percentage of de-methylated CpGs during chondrogenesis in chondrocyte chromatin states. (F) Luciferase reporter assay with enhancer regions with and without DNA methylation (n = 6). Enhancers are labelled with their nearest gene.

### SOX9 binding at chondrocyte enhancers

Transcription factor binding occurs at gene enhancers [30,31]. Therefore, we determined whether chondrocyte enhancers defined in this study contained any transcription factor binding motifs. *De novo* motif analysis using MEME-ChIP [32] revealed a SOX9 motif in chondrocyte enhancers (Fig. 5A). SOX9 is a pivotal transcription factor driving chondrogenesis and interacts with promoters and enhancers to promote chondrogenesis [33]. To further characterise chondrocyte enhancers, they were classified into two groups: new enhancers, defined by a change in chromatin state from quiescent or repressed to active enhancers during chondrogenesis, and constant enhancers; regions which were active enhancers both prior and post chondrogenesis. The analysis of motif enrichment (AME) algorithm implemented in the MEME suite of motif searching tools was used to contrast relative SOX9 motif enrichment in these two classes of enhancers found in chondrocytes. We found that both SOX9 homodimer and monomer motifs were significantly more enriched in the new enhancer class compared to the constant enhancer class (Supplementary Fig. S17). This suggests that enhancers have different properties dependent on whether they acquired enhancer status upon differentiation or if they were enhancers beforehand. To confirm whether SOX9 binds to motifs found in chondrocyte enhancers we used a publicly available mouse rib chondrocyte SOX9 ChIP-seq dataset and converted the data to human genome co-ordinates. The majority of SOX9 peaks were found in the chondrocyte strong promoter (2_TssS) state, strong active enhancer state (13_EnhS) state and quiescent state (16_Quies); the latter simply being due to the high percentage of the genome marked quiescent. Accounting for the size of chromatin states, there was more SOX9 enrichment in promoter and enhancer states (Fig. 5B). This confirms that the chondrocyte promoters and enhancers identified in our study contain real and conserved SOX9 binding sites. The impact of SOX9 overexpression was assessed on the previously cloned enhancer regions and a SOX9-responsive Col2a1 enhancer reporter (Fig. 4F). Four out of six enhancers exhibited increased enhancer activity with SOX9 overexpression (Fig. 5C). All regions except one (nearest gene *TLE3*) have a SOX9 binding site in the lifted over SOX9 ChIP-seq data; as predicted, the *TLE3* region did not show increased enhancer activity upon SOX9 overexpression.

**Figure 5.**
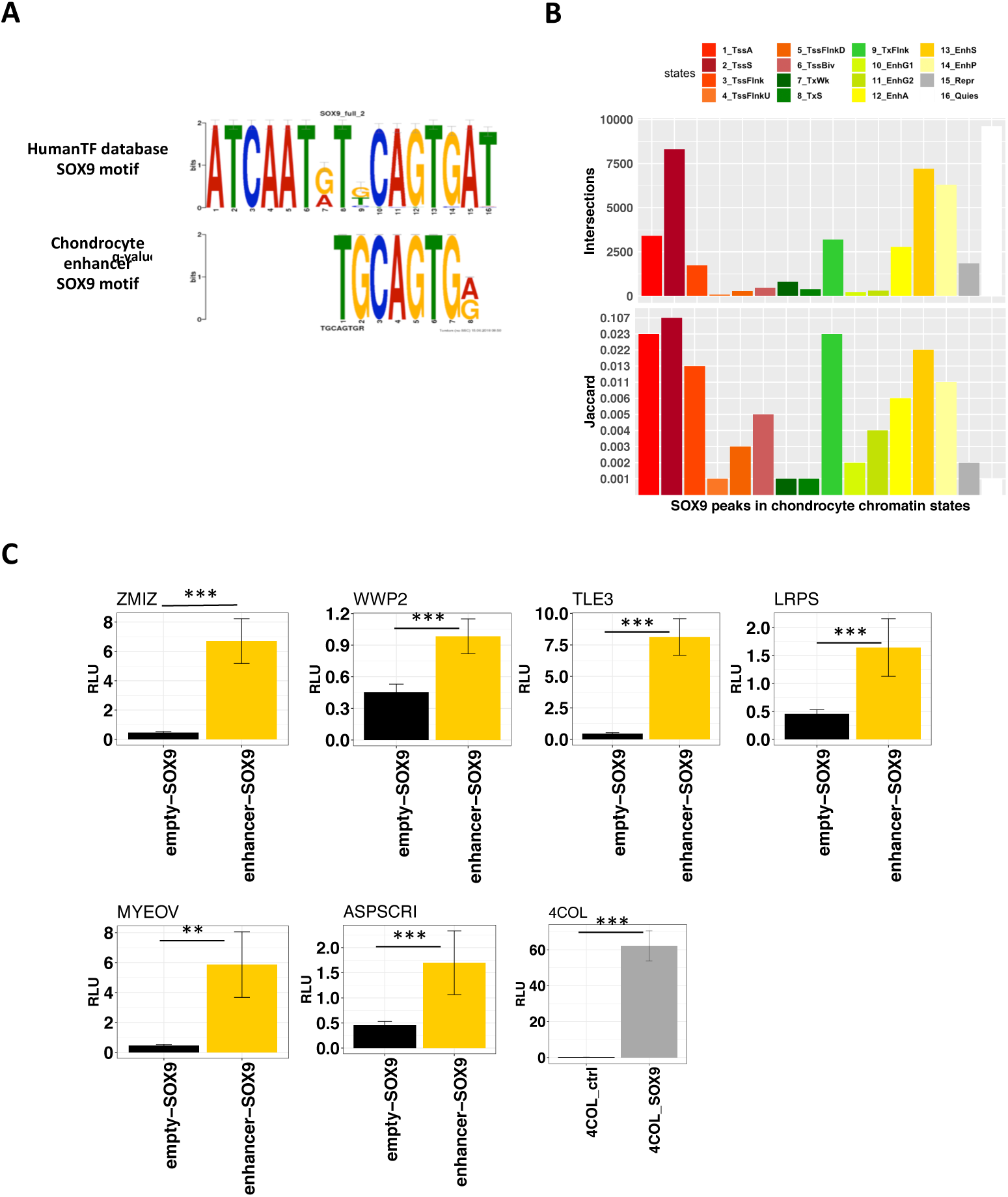
SOX9 binding in chondrocyte chromatin states. (A) SOX9 motif found in chondrocyte strong enhancer state. De novo motif analysis was performed using MEME-ChIP after randomly subsampling 10,000 individual chondrocyte enhancer states. (B) Numbers of SOX9 peaks derived from mouse rib chondrocyte ChIP-seq data in chondrocyte chromatin states. Jaccard index of similarity between SOX9 peaks and chondrocyte chromatin states. (C) Luciferase reporter assay with enhancer regions with and without SOX9 overexpression (n = 6). Enhancers are labelled with their nearest gene.

## Discussion

There are numerous *in vitro* models of chondrogenesis and although some models utilize scaffolds for cells to grow, scaffold-free models are reported to better reflect the conditions during *in vivo* chondrogenesis during development [34]. Chondrocytes from this study were derived from a scaffold-free chondrogenesis model using bone marrow derived hMSCs. Other scaffold-free models include the micromass and pellet culture system. In contrast, chondrocytes from the Roadmap project were derived from human BM-MSCs in a 3D alginate chondrogenesis model [8]. Whilst there have been some gene expression comparison between models [35,36], no comparison has been made about changes in their epigenetic landscape. Here, we show that chondrocyte gene enhancers across two different models are more concordant relative to other cell-types. This is indicative of a unique chondrocyte epigenetic signature, independent of model and laboratory specific effects. Although hMSC-derived chondrocyte enhancer concordance is evidence that chondrogenic models are reliable and comparable, their likeness to real human articular chondrocyte enhancers has yet to be investigated.

Combinations of histone modifications can define regulatory elements and regulate genes through modulating chromatin remodelling to allow or block access of transcription factors. However, histone modifications also rely on other epigenetic mechanisms such as DNA methylation and vice versa [26]. Crosstalk between the two epigenetic mechanisms allows for greater control of gene transcription and it is important to consider histone modifications in the wider context of the whole epigenome. Traditionally, studies into DNA methylation focused on gene promoters where CpG islands are more likely to be found and array probe design is biased towards promoters. Although our data is extensive, we only compared ∼450,000 (1.6%) of the ∼28 million CpG sites in the human genome [37]. Reduced representation bisulfite sequencing (RRBS) in chondrogenesis only identified limited CpG methylation changes in gene promoters [38]. However, RRBS is heavily biased towards promoters and whole genome bisulfite sequencing remains the only method that can universally capture almost the entire DNA methylome. We show in this study that significant changes occur at distal gene enhancers during chondrogenesis. DNA de-methylation at enhancer regions has also been observed during other stem cell differentiation processes, including differentiation of intestinal epithelium progenitors [39], haematopoietic stem cells [40] and embryonic stem cells [41] but also due to MSC age and culture conditions [42]. DNA de-methylation at enhancers is associated with development of most human organs [43]. Aberrant DNA methylation in enhancers has been implicated in diseases such as cancer [44–46] and osteoarthritis (OA) [47,48].

*SOX9* has a highly conserved DNA binding motif and function [49,50]. Therefore, we considered liftover of mouse reads to human genome co-ordinates to be appropriate for our analysis. Using this method, species specific SOX9 binding information is lost but conserved sites are retained, these sites arguably being the most important, as evolutionary conservation is a marker of essentiality. We found SOX9 motifs in our chondrocyte enhancers via *de novo* motif searching as well as conserved SOX9 binding using mouse SOX9 ChIP-seq. SOX9 acts in conjunction with transcription factors SOX5 and SOX6 in chondrogenesis [51], to bind to super enhancers promoting chondrogenesis [52]. Super enhancers are loosely defined as multiple enhancers in close proximity exhibiting high levels of active enhancer markers such as H3K27ac or transcription factors (Pott and Lieb, 2015). SOX9 bound enhancers have previously been proposed to be important for defining the chondrocyte phenotype. Furthermore, mutations of Sox9 binding motifs within distal *Acan* enhancers in transgenic mice resulted in a loss of chondrocyte specific expression [53].

Enhancers are thought to regulate their target genes by forming a loop to physically contact the gene promoter within topologically associating domains [54,55]; an interaction mediated by transcription factors [56]. Gene enhancers can be located distal from their target promoters and therefore target gene prediction can be challenging without chromatin conformation data [57]. Although we have validated that enhancers identified in this study do indeed possess enhancer activity that may be modulated by DNA methylation and SOX9 binding, further functional work is required to elucidate their gene target(s) and importance in cartilage development. In this study, we show that enhancers are dynamic during chondrogenesis and may serve as potential targets for modulating hMSC differentiation.

To conclude, integration of ChIP-seq with methylation data revealed that gene enhancers are de-methylated during an *in vitro* transwell model of chondrogenesis. Comparison of chromatin states across hMSCs and chondrocytes generated in this study along with those from the Roadmap Epigenomics project revealed that enhancers are more variable between cell-types compared to other chromatin states. Chondrocytes from the Epigenomics Roadmap project and this study showed a more similarity of enhancers with each other than other cell-types despite being from different models. We have established that chondrocyte enhancers contain motifs to which that SOX9 binds *in vivo*. Additional investigations are needed to elucidate further the epigenetic landscape of chondrocytes originating from other *in vitro* models and to determine whether these are comparable to the epigenome of human articular chondrocytes. A link between reactivation of developmental pathways and OA has been suggested [15]; more research is needed to fully explore the association between development and disease.

## Experimental Procedures

### hMSC Culture and Chondrogenesis

Bone marrow aspirates (donor n = 2, female, ages 22 & 24) were purchased from LONZA and hMSCs were isolated by adherence to tissue culture flasks for 24 hours. hMSCs were phenotyped by flow cytometry [58] and confirmed to have osteoblastogenic and adipogenetic potential as well as chondrogenic. Stem cells were cultured and differentiated into chondrocytes as previously described [19].

### Isolation of Chondrocytes from Cartilage-like Disc

Cartilage discs were digested at day 14 of chondrogenesis, a time point at which chondrocytes have been determined to be fully differentiated in a pellet model of chondrogenesis [59]. Cartilage discs were digested first with 1.5ml hyaluronidase (1mg/ml in sterile PBS) for 15mins at 37°C then with 1.5ml trypsin (2.5mg/ml in sterile PBS) at 37°C for 30mins. The discs were finally digested with collagenase (2mg/ml in DMEM media) for 1-1.5hrs at 37°C until fully digested and matrix was no longer visible. The digested cartilage containing media was passed through a 100μm cell strainer to remove any remaining matrix.

### Chromatin Extraction and ChIP-seq

hMSCs were harvested from monolayer culture using trypsin. Chromatin from hMSCs and differentiated chondrocytes were extracted using the Diagenode iDeal histone ChIP-seq kit (Diagenode SA, Ougrée, Belgium). Extracted chromatin was sonicated using a Diagenode Bioruptor Standard or Bioruptor Pico to an average size of 200-500bp, using 15 sonication cycles (30s on/30s off). ChIP-seq grade premium antibodies were purchased from Diagenode: H3K4me3 (included in the Diagenode iDeal histone ChIP-seq kit), H3K4me1 (Cat. no. C15410194), H3K27ac (Cat. no. C15410196), H3K27me3 (Cat. no. C15410195) and H3K36me3 (Cat. no. C15410192). Chromatin immunoprecipitation was performed following the Diagenode iDeal histone ChIP-seq protocol using chromatin from 1 million cells and 1μg antibody per ChIP. Immunoprecipitated DNA was purified using Agencourt AMPure XP beads (Beckman Coulter (UK) Ltd, High Wycombe, UK). For one hMSC chondrogenesis replicate ChIP-seq, DNA sequencing libraries were generated using Diagenode MicroPLEX v2 kit and single ended reads of 50bp length were generated on an Illumina HiSeq 2500 (Illumina Inc., San Diego, USA). The second experimental replicate was prepared using the NEBNext Ultra II kit (New England Biolabs, Hitchin, UK) and sequenced using an Illumina NextSeq 500 platform, generating 75bp single end reads.

### Luciferase Reporter Assays

Putative enhancer regions were amplified from human genomic DNA using the primers listed in Supplemental Table 2 and cloned into the pCpGL-EF1 plasmid. This plasmid has been modified from the CpG-free pCpGL-basic luciferase plasmid by addition of the EF1 CpG-free promoter upstream of the luciferase gene [60], and can thus be used to analyse DNA methylation effects on non-promoter regulatory regions. Plasmids were transformed into GT115 *E.coli* (Invitrogen) and DNA isolated using PureYield Plasmid Midiprep system (Promega). Plasmid DNA was *in vitro* methylated using CpG Methyltransferase (*M.SssI*, New England Biolabs), with the efficiency of methylation assessed by digestion using *HpaII* and *HhaI* methylation-sensitive restriction enzymes (NEB). The effect of SOX9 on enhancer activity was assessed by transfection with a SOX9 overexpression plasmid (pUT-FLAG-SOX9) [61]. A luciferase reporter (4COL) containing 4 copies of the Col2a1 48-bp enhancer was used to confirm SOX9 overexpression [62]. SW1353 chondrosarcoma cells were seeded at a cell density of 5×10^3^ per well in 96-well plates. After 24h, cells were co-transfected with 100ng (DNA methylation) or 50ng (*SOX9* overexpression) of the relevant pCpGL-EF1 plasmid and 6ng (DNA methylation) or 1.5ng (*SOX9* overexpression) of pRL-TK Renilla control plasmid and 0.3μl FuGeneHD reagent (Promega) per well. Cells were lysed 24hrs post transfection and luciferase and renilla activity measured using the Dual-Luciferase Reporter Assay kit on a Glomax-Multi reader. Three experiments with six replicates each were performed for each construct, and *luciferase*/*renilla* activity normalized to for the empty the pCpGL-EF1vector.

### ChIP-seq analysis and chromatin state learning

Quality control of sequencing reads was performed using FastQC (v.0.11.5). All reads passed quality thresholds. Reads were aligned to the reference human genome hg38 using Bowtie2 (v.2.2.4) [63]. MACS2 (v.2.1.0.2) [64] was used to call broad peaks (parameters *–broad* and *– no-model*) using input samples as controls. The ngs.plot program (v.2.61) [65] was used to visualise peak enrichment across the genome and at gene expression levels. An Illumina whole-genome expression array Human HT-12 V4 was used to determine gene expression levels prior and post chondrogenesis [19]. Normalised gene expression signals were categorised into low (signal < 7; 1^st^ quarter), medium (signal between 7 & 9) or high expression (signal > 9; 3^rd^ quarter).

ChromHMM (v.1.12) [66] was used to train a 16 state model on all histone marks assayed. The number of states was arrived at by running the model with different numbers of states until a separation of chromatin states was seen; as described by the Roadmap Epigenomics Project [22]. The Integrative Genomics Viewer (IGV) was used to visualise chromatin state tracks [67]. Global chromatin state changes between hMSC and differentiated chondrocytes were visualised using the riverplot package in R. Gene ontology (GO) terms for chromatin states were found using the GREAT tool with default settings [68]

Mouse SOX9 ChIP-seq data (GEO GSE69109) was aligned to mm10 using Bowtie2 (default settings). Aligned reads were converted to hg38 using the UCSC liftOver tool and narrow peaks were called using MACS2 (v.2.1.0.2) using input samples as control.

### Chromatin state comparisons with Roadmap Epigenomics cell-types

Chromatin state co-ordinates from our study were converted to hg19 using UCSC liftOver as Roadmap data were aligned to hg19. Similarity analysis between equivalent chromatin states across hMSC, chondrocyte and Roadmap cell-types was performed using the Jaccard index and hierarchical clustering. Roadmap chromatin state data is available to download from the project website (http://www.roadmapepigenomics.org/).

### Integration with DNA methylation

An Infinium HumanMethylation450 BeadChip array was used to quantify DNA methylation in the transwell model of chondrogenesis [27], GEO dataset GSE129266. CpG probes from the 450k methylation array were based on human reference genome hg19, therefore, CpG co-ordinates from the array were first converted to hg38 and intersected with chromatin state co-ordinates from hMSC and differentiated chondrocytes. A Chi-square test with 1000 Monte Carlo permutations was used to test independence of de-methylated CpG distribution in enhancers. All plots were generated using the ggplot2 package in R.

### Motif Analysis

A random sample of 10,000 chondrocyte enhancers identified by ChromHMM was selected for *de novo* motif analysis. MEME-ChIP (default settings) from the MEME suite of tools (Ma et al, 2014) was used to identify motifs in chondrocyte enhancers. The analysis of motif enrichment (AME) tool within MEME was used to assess relative enrichment of SOX9 binding motifs found in the footprintDB database [69] in new chondrocyte enhancers compared to constant enhancers.

## Data availability

ChIP-seq data has been deposited GSE129031. The chondrogenesis 450k DNA methylation array data can be found GSE129266. The chondrogenesis transcriptome analysis using Illumina whole-genome expression array Human HT-12 V4 is available upon reasonable request from the authors.

## Author Contributions

Kathleen Cheung: Collection and/or assembly of data, data analysis and interpretation, manuscript writing

Matthew J. Barter: Collection and/or assembly of data, provision of study material or patients

Julia Falk: Collection and/or assembly of data Carole Proctor: Conception and design

Louise N. Reynard: Conception and design, provision of study material or patients

David. A. Young: Conception and design, final approval of manuscript

## Acknowledgements

This work was supported by the Medical Research Council and Arthritis Research UK as part of the MRC-Arthritis Research UK Centre for Integrated Research into Musculoskeletal Ageing (CIMA, grant references JXR 10641 and MR/P020941/1); Arthritis Research UK [grant numbers 18746 and 19424]; the JGW Patterson Foundation; The Dunhill Medical Trust; and the NIHR Newcastle Biomedical Research.

## Data availability statement

ChIP-seq data have been submitted into the NCBI GEO data repository with accession GSE129031.

## Tables

**Supplementary table 1** – genes expressed in chondrocytes characterised into high, medium and low expression groups.

**Supplementary table 2** – Primer sequences used for cloning into luciferase enhancer constructs

## Supplementary figures

***Figure S1*** *- GREAT GO terms for chondrocyte chromatin state 1_TssA*

***Figure S2*** *- GREAT GO terms for chondrocyte chromatin state 2_TssS*

***Figure S3*** *- GREAT GO terms for chondrocyte chromatin state 3_TssFlnk*

***Figure S4*** *- GREAT GO terms for chondrocyte chromatin state 4_TssFlnkU*

***Figure S5*** *- GREAT GO terms for chondrocyte chromatin state 5_TssFlnkD*

***Figure S6*** *- GREAT GO terms for chondrocyte chromatin state 6_TssBiv*

***Figure S7*** *- GREAT GO terms for chondrocyte chromatin state 7_TxWk*

***Figure S8*** *- GREAT GO terms for chondrocyte chromatin state 8_TxS*

***Figure S9*** *- GREAT GO terms for chondrocyte chromatin state 9_TxFlnk*

***Figure S10*** *- GREAT GO terms for chondrocyte chromatin state 10_EnhG1*

***Figure S11*** *- GREAT GO terms for chondrocyte chromatin state 11_EnhG1*

***Figure S12*** *- GREAT GO terms for chondrocyte chromatin state 12_EnhA*

***Figure S13*** *- GREAT GO terms for chondrocyte chromatin state 14_EnhP*

***Figure S14*** *- GREAT GO terms for chondrocyte chromatin state 15_Repr*

***Figure S15*** *- GREAT GO terms for chondrocyte chromatin state 16_Quies*

**Figure S16** - *ChromHMM emission parameters for Roadmap’s 18 state model (A) and (B) chondrogenesis 16 state model. Roadmap’s 1_TssA state and the chondrogenesis 2_TssS state comprises equal probabilities of H3K4me3 and H3K27ac histone marks. Likewise, Roadmap’s 9_EnhA1 and chondrogenesis 13_EnhS have similar levels of H3K4me1 and H3K27ac. Other states considered comparable were Roadmap’s 5_Tx with chondrogenesis 8_TxS, Roadmap’s 6_TxWk with chondrogenesis 7_TxWk, Roadmap’s 11_EnhWk with chondrogenesis 14_EnhP, Roadmap’s 16_ReprPC with chondrogenesis 15_Repr and Roadmap’s 18_Quies with chondrogenesis 16_Quies*

**Figure S17** – *Relative enrichment of SOX9 binding motifs in new enhancers to constant enhancers. Constant enhancers were defined as enhancers that were categorized as enhancers both prior and post chondrogenesis. New enhancers became enhancers upon hMSC differentiation. The footprintDB database was searched for all SOX9 motifs, both dimer and monomer. The AME tools within MEME was used to determine relative enrichment of new enhancers compared to constant enhancers.*

